# RNAsamba: coding potential assessment using ORF and whole transcript sequence information

**DOI:** 10.1101/620880

**Authors:** Antonio P. Camargo, Vsevolod Sourkov, Marcelo F. Carazzolle

**Affiliations:** Department of Genetics, Evolution, Microbiology and Immunology, Institute of Biology, University of Campinas, Campinas, SP, 13083-862, Brazil; Graduate Program in Genetics and Molecular Biology, Institute of Biology, University of Campinas, Campinas, SP, 13083-862, Brazil; Department of Computer Science, ReDNA Labs.

## Abstract

**Motivation:** The advent of high-throughput sequencing technologies made it possible to obtain large volumes of genetic information, quickly and inexpensively. Thus, many efforts are devoted to unveil the biological roles of genomic elements, being one of the main tasks the identification of protein-coding and long non-coding RNAs.

**Results:** We describe RNAsamba, a tool to predict the coding potential of RNA molecules from sequence information using a deep-learning model that processes both the whole sequence and the ORF to look for patterns that distinguish coding and non-coding RNAs. We evaluated the model in the classification of coding and non-coding transcripts of humans and five other model organisms and show that RNAsamba mostly outperforms other state-of-the-art methods. We also show that RNAsamba can identify coding signals in partial-length ORFs and UTR sequences, evidencing that its model is not dependent on the presence of complete coding regions. RNAsamba is a fast and easy tool that can provide valuable contributions to genome annotation pipelines.

**Availability and implementation:** The source code of RNAsamba is freely available at:https://github.com/apcamargo/RNAsamba.

## 1. Introduction

High-throughput sequencing technology has enabled the sequencing of genomes and transcriptomes of a myriad of species, yielding large quantities of genetic information [1]. Hence, great effort is dedicated to characterize the obtained data, mainly by the identification of functional genomic elements such as messenger RNAs (mRNAs) and long non-coding RNAs (lncRNAs).

Due to their role of carriers of protein synthesis information, mRNAs have been studied for several decades and are well represented in genetic databases. In contrast, lncRNAs, which are defined as transcripts longer than 200 nucleotides that are not translated into proteins [2], have been known for much less time and only recently their role as regulators of gene expression and their link to genetic diseases have been unveiled.

One of the main goals of the functional annotation of genomes and transcriptomes is the identification of mRNAs and lncRNAs. Usually, this process relies on the comparison of sequences or structures with databases of biological sequences, which is very time-consuming [3] and poses limitations for both the annotation of mRNAs and lncRNAs. As only a fraction of the genetic diversity existing in nature is known and available in databases, many new protein-coding genes are not identified as such because their protein product is not found among existing data. On the other side, as lncRNAs are not under the same evolutionary constraints as mRNAs, they display lower sequence conservation than protein-coding transcripts [4, 5], resulting in failure to find homologous sequences in database searches [6, 7].

Even though mRNAs and lncRNAs usually share many molecular features [8, 9], they display contrasting sequence characteristics that can be used to create statistical models that can compute the coding potential of any given transcript without the limitations of database-based annotation pipelines. Most of these approaches employ machine learning algorithms to differentiate coding and non-coding transcripts based on a series of human-designed sequence features such as ORF length and integrity [10, 11], GC-content [8], 3-base periodicity [12], k-mer frequencies [13, 14] and hexamer usage bias [15]. The usage of these features, however, may introduce bias to the classification, causing, for instance, the models to misclassify lncRNAs possessing long ORFs and coding transcripts containing short or truncated ORFs.

The power of multi-layered neural networks to identify deep patterns has made them the de facto standard in many machine learning applications, such as image and text analysis, and have been extensively employed in bioinformatics to provide new biological insights [16]. Contrasting to conventional machine learning algorithms, deep learning approaches do not rely on human-designed features and can be used to capture concealed sequence signals that are fundamentally different between mRNAs and lncRNAs.

Here we describe RNAsamba, a tool that uses a novel neural network architecture to tackle the mRNA/lncRNA classification problem relying solely on sequence information. We show that our method outperforms previous tools in a variety of metrics and is robust to limitations commonly found in real world data, such as truncated ORFs.

## 2. Background

Recurrent Neural Networks (RNNs) are a type of neural network in which each node takes the output of a previous node as input, forming a directed graph. This architecture confers RNNs the property of remembering previous states, making them ideal to deal with sequential data such as nucleotide sequences [16]. One well documented drawback of traditional RNNs is the issue of long-range dependencies, which hinders the training of networks with sequences longer than a few hundred time steps and makes it difficult to train RNNs with long sequences [17]. To tackle this problem, the recently introduced IGLOO [18] architecture looks at sequences as a whole, rather than sequentially like in the recurrent paradigm. To do so, IGLOO creates representations of sequences by taking patches of the feature space and multiplying them by learnable weights (Figure 1A).

**Figure 1.**
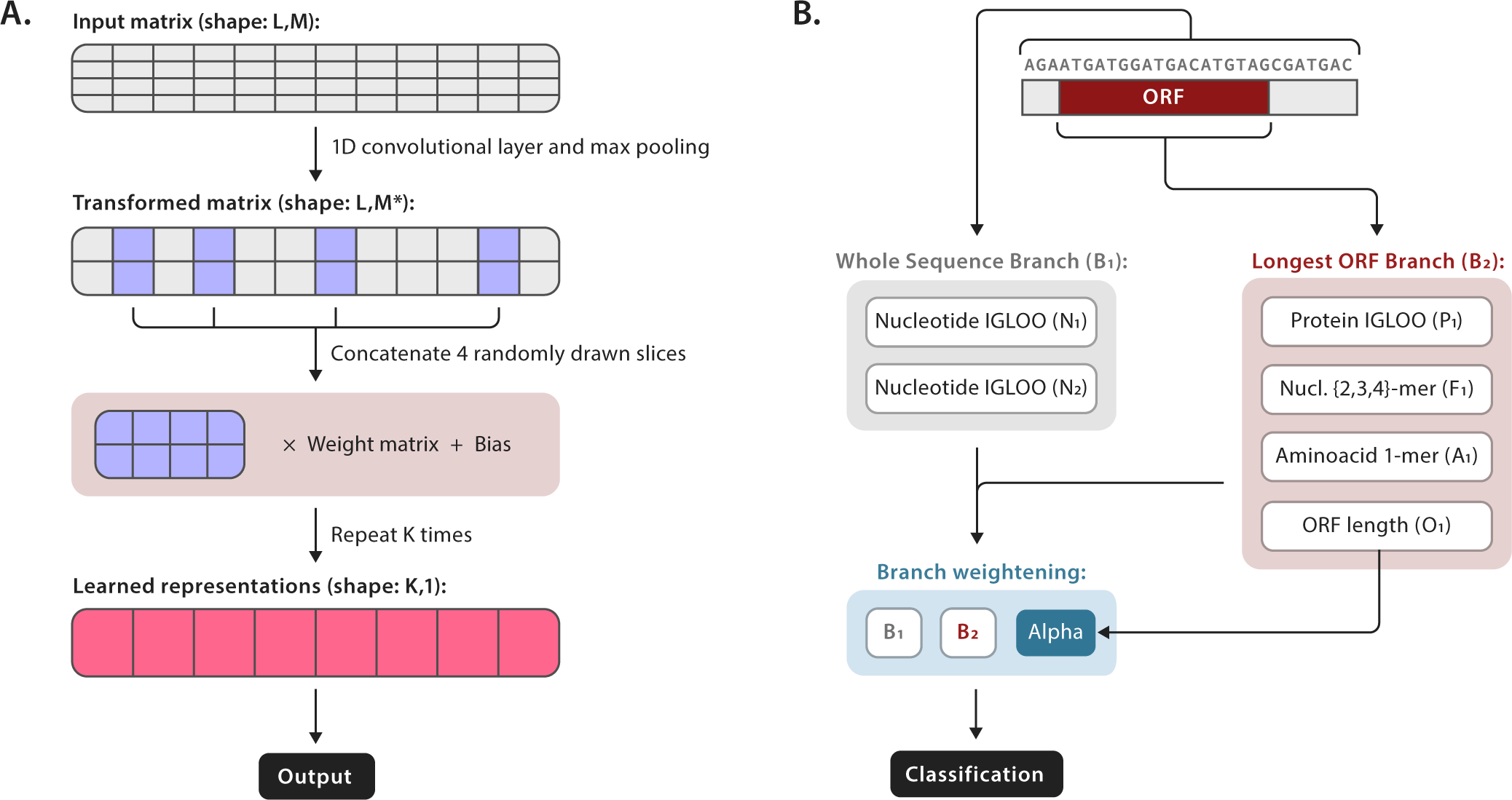
(A) In an IGLOO layer the input sequence is initially processed by an 1D convolutional layer and down-sampled using the max pooling approach. From the resulting matrix, K patches consisting of four random slices are drawn from the matrix and multiplied by K learnable weights, producing a high-level representation of the sequence input. (B) From the RNA sequence RNAsamba derives two branches. In the Whole Sequence Branch (B_1_), the whole transcript nucleotide sequence is fed to two IGLOO layers to create high-level representations of the transcript (N_1_ and N_2_). In the Longest ORF Branch (B_2_) four layers are derived from the ORF sequence: an IGLOO representation of the putative protein (P_1_), nucleotide k-mer frequencies (F_1_), aminoacid frequencies (A_1_) and the ORF length (O_1_). The two branches are weighted by the α parameter and then used to compute the final classification of the transcript.

In an IGLOO layer, input sequences are of shape (L,M), where L is the length of the sequence and M is feature size, i.e. the size of the representation of the element at a given position. IGLOO uses an initial 1-D convolutional layer and max pooling to transform the input into a (L,M*)-shaped array, which can be scaled to accommodate for the overall size of the network. Then, IGLOO iteratively collects K patches, each containing 4 random matrix slices, which are multiplied by learnable weights and joined, resulting in a K-sized representation of the sequence. Intuitively, the weight learns relationships between non-necessarily contiguous slices of the feature map. Using K of those weights allows the network to find a sequence representation composed of K different non-local relationships. This representation can then be fed to a dense layer for classification.

By taking global snapshots of the sequence, IGLOO networks can be used to process very long sequences, making them particularly interesting for nucleotide sequence data. Furthermore, IGLOO layers can be easily parallelized and run significantly faster than RNN variants, such as GRUs and LSTMs, for a similar number of trainable parameters.

## 3. Algorithm

Starting from the initial nucleotide sequence, RNAsamba computes the coding potential of a given transcript by combining information coming from two different sources (Figure 1B): the Whole Sequence Branch (B_1_) and the Longest ORF Branch (B_2_). B_1_ contains whole sequence representations of the transcript and can capture protein-coding signatures irrespective of the identification of the ORF. In contrast, B_2_ carries information extracted from the longest identified ORF and the putative protein translated from it. By taking into account these two sources of sequence information, RNAsamba builds a thorough model of the transcript, improving the classification performance of the algorithm.

### ORF extraction

RNAsamba scans each of the three reading frames looking for fragments that initiate with a start codon (`ATG`) and finish either with a stop codon (`TAG, TAA` or `TGA`) or at the end of the transcript. The longest fragment among the ones found in all reading frames is then extracted, regardless of finishing with a stop codon or not.

### Sequence processing and encoding

RNAsamba generates high-level representations of both nucleotide and aminoacid sequences using IGLOO units. As these units require fixed length sequences as input, transcript and protein sequences are truncated to a maximum length of 3,000 nucleotides and 1,000 aminoacids, respectively. Even though these thresholds were arbitrarily chosen, we observed that, while using them, the algorithm exhibits faster training times and can capture enough information to correctly classify very long transcripts (Table 1). We believe that this because the region that contributes the most to classification is located right after the start codon [19]. The sequences are then converted into numeric representations as follows:

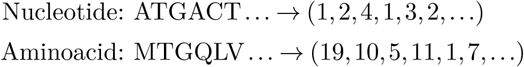

Finally, nucleotide and protein sequences shorter than the maximum length threshold are then zero-padded to 3,000 and 1,000 elements, respectively.

**Table 1.** Ablation studies of the RNAsamba model. Default parameters are highlighted in bold. Reported train times, test times and accuracy values correspond to the average of five independent executions. Computations were performed with two Intel(R) Xeon(R) E5-2420 v2 CPUs.

| Architecture | Maximum length (nt/aa) | Branches | Training (s) | Inference (s) | Accuracy |
| --- | --- | --- | --- | --- | --- |
| **IGLOO** | **3000/1000** | B<sub>1</sub>, B<sub>2</sub> | **965.41** | **94.74** | **0.9393** |
| IGLOO | 2400/800 | B<sub>1</sub>, B<sub>2</sub> | 918.13 | 93.17 | 0.9319 |
| IGLOO | 4500/1500 | B<sub>1</sub>, B<sub>2</sub> | 1110.37 | 98.38 | 0.9390 |
| IGLOO | 6000/2000 | B<sub>1</sub>, B<sub>2</sub> | 1242.86 | 103.04 | 0.9392 |
| IGLOO | 3000/1000 | B<sub>1</sub> | 658.83 | 28.62 | 0.7814 |
| GRU | 3000/1000 | B<sub>1</sub>, B<sub>2</sub> | 8853.15 | 306.34 | 0.9110 |
| LSTM | 3000/1000 | B<sub>1</sub>, B<sub>2</sub> | 10468.63 | 510.63 | 0.9089 |

### Whole Sequence Branch (B_1_)

To obtain high-level representations of the transcript, the whole nucleotide sequence is inputted into two independent stacked IGLOO units, N_1_ and N_2_, with K_1_ (K_1_ = 900) patches and distinct kernel sizes in their initial convolutional layers. The outputs of these units are then concatenated and fed to a dense layer resulting in B_1_.

### Longest ORF Branch (B_2_)

B_2_ is the result of the combination of four different layers that carry different properties of the ORF sequence. Layer P_1_ contains a representation of the protein sequence and is obtained by inputting the aminoacid sequence of the putative protein into an stacked IGLOO layer with K_2_ (K_2_ = 600) patches; layer F_1_ is comprised of the relative frequencies of nucleotide k-mers (*k* ⊂ {2, 3, 4}) in the ORF; layer A_1_ contains the relative aminoacid frequency of the translated ORF; layer O_1_ consists of the length of the longest identified ORF. B_2_ is obtained by feeding P_1_, F_1_, A_1_ and O_1_ to four independent dense layers, concatenating the outputs into a single matrix that is then fed a final dense layer.

### Branch weightening

The branches B_1_ and B_2_ gather different information from the transcript: while B_1_ captures patterns from the whole transcript sequence, B_2_ picks up information specific to the ORF. Therefore, we include an attention mechanism, the α parameter, to weight information coming from these two branches. This mechanism is important, for instance, to correctly classify transcripts with unusual ORF length, such as noncoding transcripts with long ORFs or truncated protein-coding RNAs.

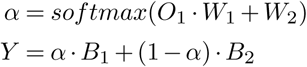

Where W_1_ and W_2_ are trainable matrices and α is a matrix that is used to weight B_1_ and B_2_ in the final layer (Y). While training the algorithm end-to-end, the weights in W_1_ and W_2_ are optimized to maximize classification accuracy.

To obtain the coding score, Y is fed to a dense layer with a softmax activation that normalizes the input into probabilities for each class [20]. Training is performed by minimizing the categorical cross-entropy using the Adam optimizer [21].

## 4. Implementation

RNAsamba is built with popular state-of-the-art deep learning libraries, TensorFlow [22] and Keras. We provide an execution guide and convenient scripts to make the process of training new models and classifying transcripts easy for the end user. For training new models, RNAsamba supports changing the number of epochs and batch size. It also allows the user to enable early stopping, which is useful to avoid overfitting. For inference, our implementation allows the input of multiple weights files that are combined in an ensemble classification, also helping to reduce model variance.

## 5. Results

### RNAsamba can accurately distinguish mRNAs from lncRNAs in several datasets

To evaluate the ability of RNAsamba’s algorithm to learn how to discriminate coding sequences from non-coding ones, we compared it with five state-of-the-art coding potential predictors: CPAT [23], CPC2 [24], FEELnc [25], lncRNAnet [26] and mRNN [19], being the last two based on neural network models. To keep the comparison as unbiased as possible, the benchmark was performed using four independent datasets consisting of coding and non-coding human transcripts previously used in the literature. These datasets exhibit differing characteristics regarding balance, transcript, and ORF length distributions (Table S1 and Figure S1). When possible, classification models were trained using the training set of each dataset. As data augmentation is a central feature of mRNN’s train process, we chose not to train new models for this algorithm.

We found that RNAsamba outperforms the other predictors in a variety of metrics in all datasets (Figure 2A and Table S2). The only dataset in which RNAsamba does not show consistently better classification performance is the mRNN-Challenge dataset, where mRNN displays better overall results. It should be noted, however, that mRNN was tested with a pre-trained model (unfilled circles in Figure 2) made available by its developers and we did not reproduce their training procedure, which involves data augmentation of the training sequences.

**Figure 2.**
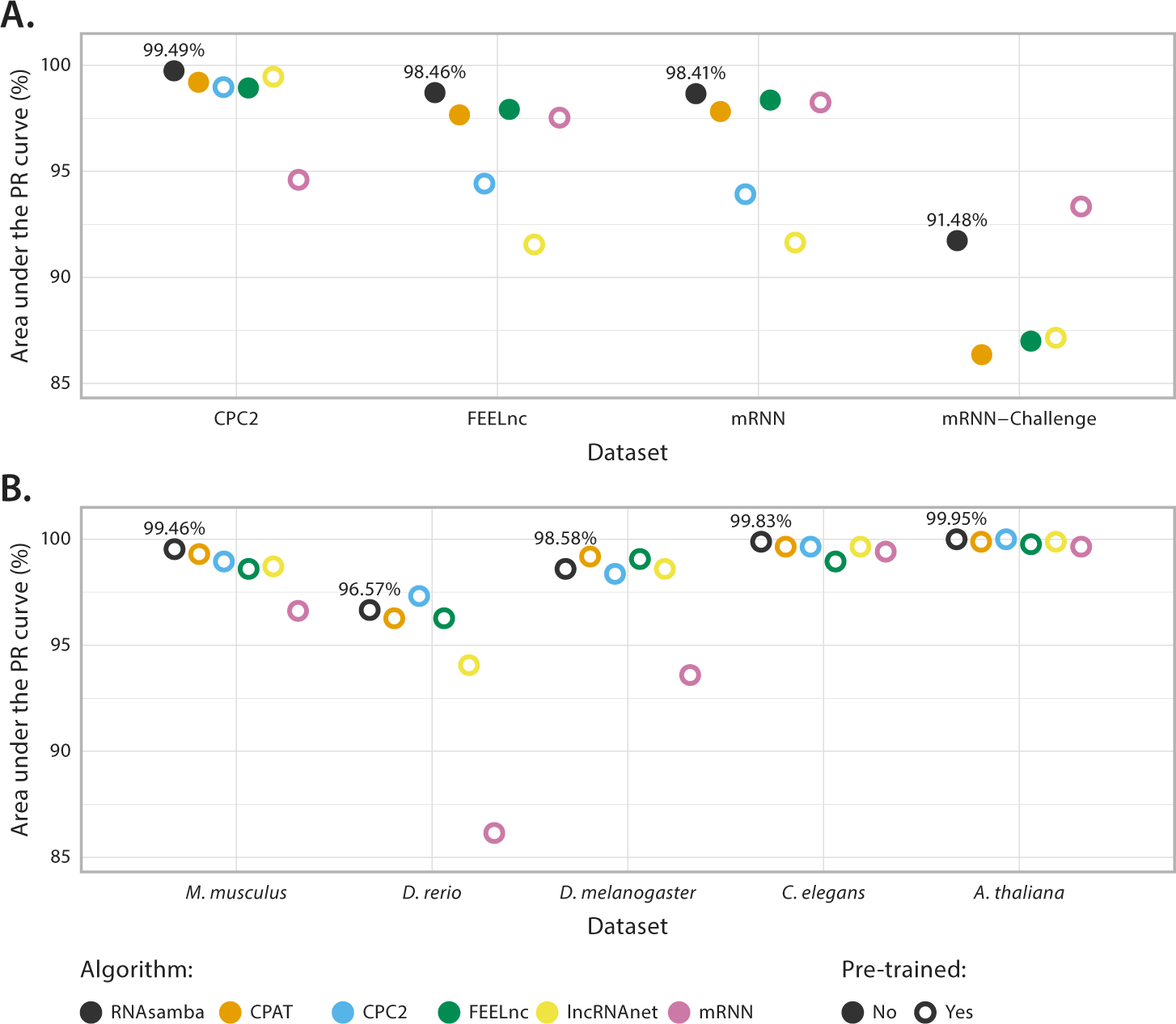
Classification benchmark of six different coding potential calculators. (A) Classifiers performance in four independent test datasets containing human transcripts. Values correspond to the area under the precision-recall curve. Models that were trained with the corresponding training sets are represented by filled circles, while pre-trained models are indicated by unfilled circles. CPC2 is outside of the displayed range in the mRNN-Challenge test dataset (75.35%). (B) Classifiers performance in five different species. Values correspond to the area under the precision-recall curve.

### RNAsamba’s model generalizes to different species

In order to evaluate if a RNAsamba model trained with human RNA sequences generalizes well to other species, we evaluated a model trained with human data in multiple test datasets, each containing both mRNAs and ncRNAs from one of five different species: *M. musculus, D. rerio, D. melanogaster, C. elegans* and *A. thaliana* (Table S3 and Figure S2). We also compared the performance of RNAsamba to five other algorithms pre-trained with human transcripts.

RNAsamba exhibits good classification performance in every species, irrespective of the evolutionary distance to humans, showing that a model learned from human sequence data can be generalized to different organisms. When compared to other tools, RNAsamba recurrently is placed among the best tools, showing slightly worse results only in *D. rerio* and *D. melanogaster*, where it displays a drop in precision (Figure 2B and Table S4). Notably, mRNN exhibits a significant decrease in classification performance when compared to its results in human data, evidencing that its algorithm may not handle well RNA sequences from different species.

### RNAsamba can identify truncated coding sequences

Since it comprehends the coding portion of the RNA, the ORF is generally used as the main source of information to detect potential protein-coding transcripts. Because of that, most mRNA predictors use human-engineered features extracted from the coding portion of the transcript, such as the ORF length and coverage. This dependence of a detectable in-frame ORF to identify coding sequences impairs the function of these algorithms to annotate the majority of transcriptome datasets, which contain a large fraction of partial-length transcripts [6, 27, 28].

As the B_1_ branch of RNAsamba captures sequence information that is independent of the ORF, it can detect protein-coding signatures even in the absence of a start codon. Thus, we tested the algorithm’s performance in the identification of truncated mRNA transcripts in which both the start and stop codon are absent. To avoid biases caused by the detection of a fragment of the true ORF, we also evaluated RNAsamba’s performance in a separate set of truncated transcripts that show no in-frame start codon inside the ORF, meaning that the model would have to capture ORF-independent coding marks to identify mRNAs. For this test, we trained RNAsamba with both complete and truncated sequences, aiming to make the algorithm more capable of identifying mRNAs by looking at the whole sequence context.

Inspection of the fraction of identified mRNAs obtained from each stratum of truncated ORFs revealed that RNAsamba is capable of identifying a substantial fraction of the mRNAs even when most of the ORF is absent (Figure 3). We also noted a negative association between the amount of available ORF information and the median value of the α parameter, showing that RNAsamba favors B_1_ as ORF-derived data becomes sparse (Figure S3).

**Figure 3.**
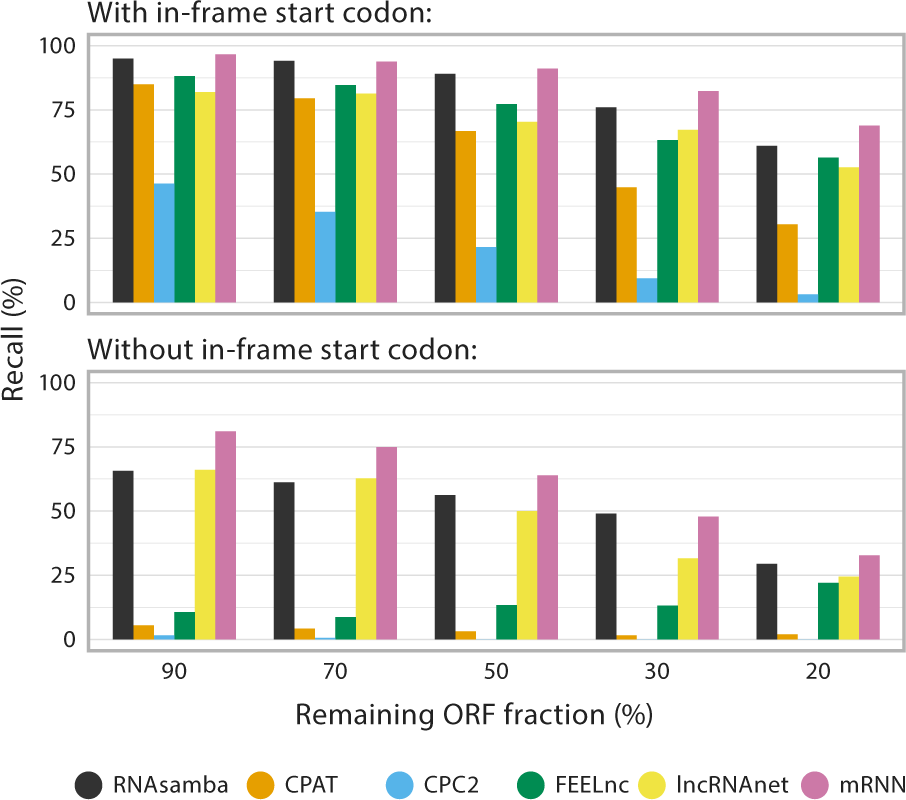
Evaluation of the ability of different tools to detect the coding potential of ORFs with varying degrees of fragmentation.

When contrasted to three ORF-dependent algorithms, CPAT, CPC2 and FEELnc, RNAsamba displayed a much better performance at identifying partial coding sequences. The discrepancy between RNAsamba and these algorithms is much more pronounced in the case of the truncated transcripts without in-frame start codons, as CPAT, CPC2 and FEELnc are incapable of finding fragments of the true ORF, making their predictions mostly unreliable. When compared to other algorithms that don’t strictly rely on ORF sequences, RNAsamba displays better classification performance than lncRNAnet, but generally worse than the mRNN model. We suspect that mRNN’s good performance in this specific kind of data is possibly due to the use of artificially introduced reading frame shifts during the data augmentation process [19].

### RNAsamba can detect a translation-related sequence residing outside of the ORF

The Kozak consensus sequence, which spawns from the −6 to the +4 positions of mRNAs, is a recurring sequence in coding transcripts [29] and plays a major role in the initiation of the translation process [30], evidencing that portions of untranslated regions can affect translation efficiency. As RNAsamba uses whole-sequence information to process RNA sequence data, we investigated whether its algorithm is sensitive to changes in the Kozak sequence region.

Thus, for each of 1,000 randomly chosen mouse mRNAs, we derived two sets containing 100 computer-generated transcripts each. In the control set, new sequences were created by exchanging the Kozak sequence region of the mRNA by fragments created by sampling nucleotides from a uniform probability distribution. In contrast, nucleotides of the computer-generated fragments of the second set were sampled according to the probability distribution of the Kozak consensus sequence (Figure S4).

We found that Kozak-derived sequences lead to an overall increase of transcripts’ coding score. In the majority of the tested transcripts (77.71%), this score was significantly larger (FDR-adjusted *p*-*value* ≤ 0.05) in fragments generated from the Kozak consensus probability distribution, indicating that RNAsamba is able to detect an important signal that contributes to mRNA translation even though it mostly resides outside of the ORF. Accordingly, we observed that there is a significant (*p*-*value ≈* 0.01) negative correlation between the coding score of a given sequence and the Hamming distance between its computed-generated portion and the Kozak sequence consensus.

We also investigated whether the effect of the Kozak sequence on the coding score is diminished in longer sequences, since they intrinsically carry larger amounts of information to be processed by the RNAsamba algorithm. We noticed that for transcripts longer than a well-defined threshold, around 3,160 base pairs (bp), there is no detectable variation among the coding scores of the control and the Kozak-derived groups (Figure S5), suggesting that the effect of this short signal is no longer detectable as the algorithm processes larger chunks of information.

### RNAsamba is faster than neural network-based alternatives

Neural networks models are becoming increasingly popular due to their ability to learn nonintuitive patterns, that would otherwise be ignored by humans, from large quantities of data. This learning power is, however, accompanied by an enormous increase in the number of trainable parameters when compared to traditional machine learning techniques, greatly increasing training time [16]. We felt that the available neural network-based coding-potential calculators impose a barrier for most users, as they do not possess GPU hardware to increase performance. By using modern libraries and IGLOO layers we sought to develop an algorithm that makes it feasible to train new models even with traditional CPUs.

We compared RNAsamba to lncRNAnet and mRNN with respect to inference and training times using the FEELnc dataset. These two algorithms employ traditional RNN variations — LSTM in lncRNAnet and GRU in mRNN — that were previously shown to be slower than IGLOO [18]. Indeed, we found that RNAsamba’s inference is, on average, 10.5 and 3.6 times faster than lncRNAnet and mRNN, respectively (Figure 4A). Regarding training, RNAsamba is 14.2 faster than mRNN (Figure 4B). Jointly, these results show that RNAsamba is faster than current alternatives, making it more accessible to most users.

**Figure 4.**
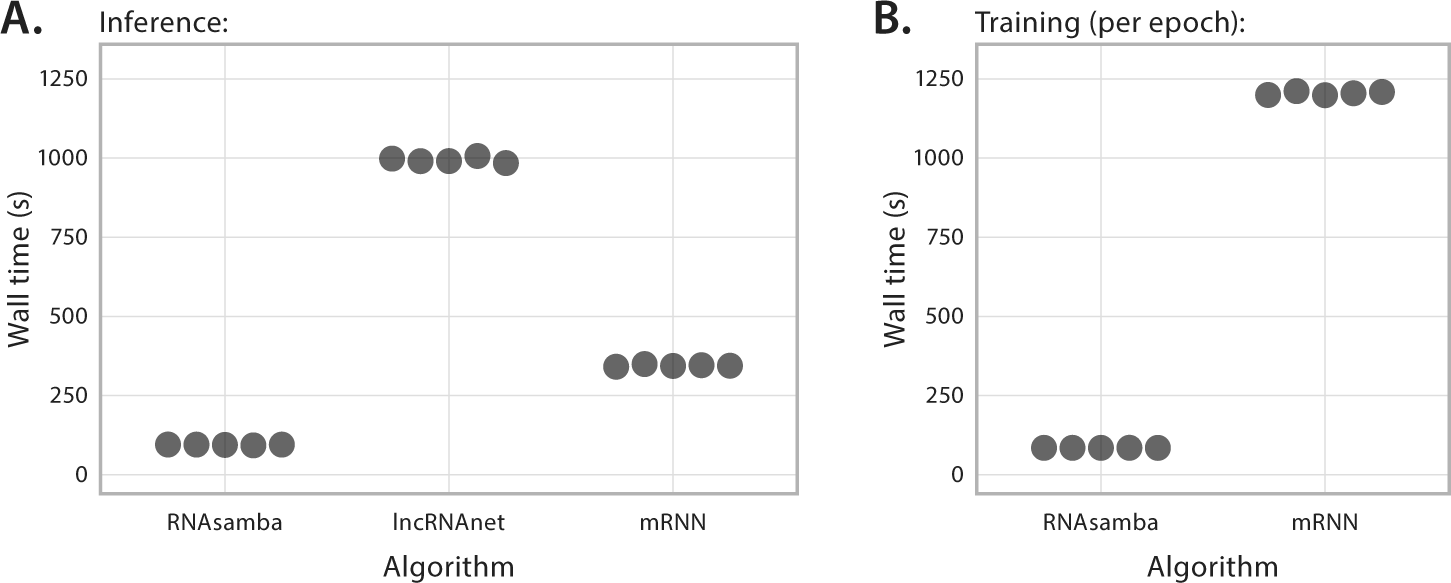
Computational performance of RNAsamba, lncRNAnet and mRNN. (A) Inference wall time of five independent executions of each algorithm. (B) Average wall time per epoch of five independent executions of RNAsamba and mRNN. LncRNAnet does not provide an interface to train new models, thus its training times were not measured. Computations were performed with two Intel(R) Xeon(R) E5-2420 v2 CPUs.

### Ablation studies

We investigated the effect of altering some of the features of RNAsamba’s algorithm to its overall performance.

#### Changing the maximum sequence length

As IGLOO layers require fixed-length inputs, we arbitrarily chose to truncate nucleotide and aminoacid sequences at the positions 3,000 and 1,000, respectively. To check whether this choice negatively affected RNAsamba’s classification performance by not providing it with important sequence information, we developed two alternative versions of the model that truncate nucleotide and aminoacid sequences at 4,500/1,500 and 6,000/2,000. We verified that raising the input sequences maximum length increased both the train and test times, without improving the model’s accuracy. Reducing the maximum lengths to 2,400/800 resulted in a slight drop in classification performance (Table 1).

#### Removing the B_2_ branch

By removing the B_2_ branch we deprived RNAsamba’s algorithm of ORF-derived features, forcing it to leverage whole-sequence information to distinguish between mRNAs and lncRNAs. We observed that this ablation reduced the accuracy of the network by 15.79% (Table 1), leading us to the conclusion that the features the algorithm derives from the ORF contain key information that is not extracted from the nucleotide sequence by the IGLOO layer alone.

#### Replacing IGLOO with GRU and LSTM

The Gated Recurrent Unit (GRU) [31] and the Long Short-Term Memory (LSTM) [32] are established RNN architectures, commonly used in deep-learning tasks that deal with sequences. Recently, IGLOO has been shown to outperform both GRU and LSTM in terms of run time and accuracy on some standard benchmark problems such as the copy-memory and the addition tasks [18]. To evaluate whether this holds true in the mRNA/lncRNA classification paradigm, we developed alternative versions of our algorithm in which IGLOO was substituted by GRU or LSTM layers with 256 units. We found that the model using IGLOO is more accurate and significantly faster, for both training and classification, than the GRU and LSTM variants (Table 1).

## 6. Conclusion

In this study, we presented RNAsamba, a new deep learning-based tool to predict the coding potential of RNA transcripts relying solely in sequence information. Compared to other algorithms, RNAsamba exhibits better classification performance in multiple human datasets and generalizes very well to other species, without relying on computationally-expansive data augmentation.

We believe that RNAsamba’s algorithm introduces two major contributions: (1) the usage of the IGLOO architecture to learn from sequence data and (2) the integration of whole transcript and ORF-derived information into a single coding score. By using IGLOO layers, RNAsamba can learn nonintuitive coding patterns, as we demonstrated with the Kozak consensus, without relying on biased human-designed features. This architecture also makes RNAsamba significantly faster than RNN-based algorithms, making it more appealing to most users. Through the usage of its two branches, RNAsamba can identify mRNAs with short or incomplete ORFs, which usually are misclassified by most algorithms.

With RNAsamba, we sought to offer a fast and easy-to-use tool to most researchers. To achieve that, we developed our software using modern and well documented libraries. Also, we provide convenient scripts to promptly execute training and inference tasks. By doing so, we believe that RNAsamba provides most users with a state-of-the-art coding potential predictor that can be easily used to accurately predict mRNAs and lncRNAs in genome annotation pipelines.

## 7. Materials and methods

### Classification performance evaluation

We assessed the performance of RNAsamba and five other sequence-dependent classification software: CPAT (1.2.4), CPC2, FEELnc (version 0.1.1), lncRNAnet and mRNN. We calculated the performance metrics considering mRNAs as the positive class and ncRNAs as the negative class.

For the evaluation in each of the human test datasets, RNAsamba, CPAT and FEELnc were trained with the corresponding train sets. We used pre-trained models for CPC2, lncRNAnet and mRNN. For the classification evaluation in the *M. musculus, D. rerio, D. melanogaster, C. elegans* and *A. thaliana* datasets, RNAsamba was trained with the sequences of all four human datasets. Other programs were executed with their pre-trained models. mRNN was loaded with weights provided in the `w14u3.pkl` file.

Links for download of the datasets used in these benchmarks can be found in the Supplementary Data.

### Truncated ORFs dataset

To generate the test for the analysis of truncated transcripts, mouse ORF sequences were retrieved from Ensembl (release 94) [33] and sequences shorter than 300 nucleotides were discarded. Next, ORFs that exhibited an in-frame start codon and the ones that didn’t were separated into different sets. The start and stop codons were removed from the sequences of both sets, guaranteeing that the true beginning and end of the ORFs would not be detected by the classifiers. Subsequently, each set was used to generate five subsets consisting of 1,000 randomly sampled sequences. Finally, the sequences from each dataset were sliced at random positions to generate sets of fragmented ORFs with fixed relative lengths (20%, 30%, 50%, 70% and 90% of the total ORF length).

For the performance evaluation, we used a RNAsamba model trained with a set containing the CPC2, FEELnc and mRNN human train and test sets as well as fragmented ORFs extracted from 50,000 of those sequences. CPAT, CPC2 FEELnc, lncRNAnet and mRNN were executed using pretrained models. mRNN was loaded with weights provided in the `w14u3.pkl` file.

### Kozak sequence analysis

The 100 different 10 bp fragments in the Kozak sequence set and the control set were generated, respectively, from the Kozak sequence probability distribution (Figure S4A) and a uniform distribution, in which all four nucleotides are equally probable to be drawn in each position (except for the start codon). The distance between the generated fragments and the Kozak sequence was obtained by computing their Hamming distances to two sequences derived from the Kozak consensus (GCC[AG]CCATGG) and choosing the lowest value.

We randomly selected 1,000 sequences among mouse mRNAs, retrieved from Ensembl (release 94), whose 5’ UTR contained at least 6 nucleotides. Then, the region spawning the positions −6 to +1 of each mRNA was exchanged by the 10 bp fragments of the Kozak sequence set and control set, producing two sets of hybrid transcripts containing both biological and computer-generated sequences (Figure S4B).

For each mRNA, we used one-tailed Mann-Whitney tests to test for differences in the coding scores of sequences in the two sets. We used the Fisher’s method to aggregate p-values and the Benjamini-Hochberg procedure to compute the false discovery rate (FDR). Kendall’s tau coefficient was used to measure the degree of association between coding scores and Hamming distance to the Kozak sequence.

### Ablation studies

The models generated in the ablation studies were trained for 10 epochs using the FEELnc human train set and all the performance evaluations were measured using the FEELnc human test set.

## Code and data availability

The source code for RNAsamba is available in an online repository (https://github.com/apcamargo/RNAsamba). Train and test sequences generated for the truncated ORF analysis, as well as computer-generated Kozak fragments were uploaded to Open Science Framework (https://doi.org/10.17605/OSF.IO/MD56Y).

