## Supplementary material for "RNAsamba: coding potential assessment using ORF and whole transcript sequence information"

Antonio P. Camargo, Vsevolod Sourkov, and Marcelo F. Carazzolle

**Table S1.** Number of transcripts belonging to each class in the test sets.

| | Coding | Non-coding |
|---|---|---|
| CPC2 | 6,142 | 12,019 |
| FEELnc | 5,000 | 5,000 |
| mRNN | 500 | 500 |
| mRNN-Challenge | 500 | 500 |

**Table S2.** Comparison of classification performance in four distinct datasets. When possible, classification models were trained using the designed training set for each dataset (●). As new models could not be trained for some of the tested algorithms, publicly available pre-trained models were used instead (○). MCC: Matthews correlation coefficient. PRC: area under the precision-recall curve

| | Pre-trained | Accuracy | Precision | Recall | MCC | F1-score | PRC |
|---|---|---|---|---|---|---|---|
| CPC2 dataset | | | | | | | |
| RNAsamba | ● | 0.9769 | 0.9618 | 0.9700 | 0.9484 | 0.9659 | 0.9949 |
| CPAT | ● | 0.9633 | 0.9279 | 0.9666 | 0.9194 | 0.9469 | 0.9895 |
| CPC2 | ○ | 0.9562 | 0.9140 | 0.9609 | 0.9041 | 0.9369 | 0.9872 |
| FEELnc | ● | 0.9605 | 0.9227 | 0.9637 | 0.9133 | 0.9429 | 0.9868 |
| lncRNAnet | ○ | 0.9768 | 0.9348 | 0.9841 | 0.9433 | 0.9588 | 0.9921 |
| mRNN | ○ | 0.9524 | 0.8988 | 0.9682 | 0.8972 | 0.9322 | 0.9435 |
| FEELnc dataset | | | | | | | |
| RNAsamba | ● | 0.9425 | 0.9654 | 0.9178 | 0.8860 | 0.9410 | 0.9846 |
| CPAT | ● | 0.9266 | 0.9468 | 0.9040 | 0.8541 | 0.9249 | 0.9741 |
| CPC2 | ○ | 0.8502 | 0.9298 | 0.7576 | 0.7127 | 0.8349 | 0.9417 |
| FEELnc | ● | 0.9229 | 0.9237 | 0.9220 | 0.8458 | 0.9228 | 0.9767 |
| lncRNAnet | ○ | 0.8689 | 0.8040 | 0.9754 | 0.7551 | 0.8815 | 0.9130 |
| mRNN | ○ | 0.9389 | 0.9412 | 0.9362 | 0.8778 | 0.9387 | 0.9728 |
| mRNN dataset | | | | | | | |
| RNAsamba | ● | 0.9340 | 0.9109 | 0.9620 | 0.8693 | 0.9358 | 0.9841 |
| CPAT | ● | 0.8720 | 0.8218 | 0.9500 | 0.7532 | 0.8813 | 0.9757 |
| CPC2 | ○ | 0.8400 | 0.9187 | 0.7460 | 0.6923 | 0.8234 | 0.9367 |
| FEELnc | ● | 0.9300 | 0.9283 | 0.9320 | 0.8600 | 0.9301 | 0.9811 |
| lncRNAnet | ○ | 0.8610 | 0.7915 | 0.9800 | 0.7433 | 0.8750 | 0.9139 |
| mRNN | ○ | 0.9510 | 0.9480 | 0.9520 | 0.9000 | 0.9500 | 0.9800 |
| mRNN-Challenge dataset | | | | | | | |
| RNAsamba | ● | 0.8340 | 0.8646 | 0.7920 | 0.6700 | 0.8260 | 0.9148 |
| CPAT | ● | 0.7300 | 0.7357 | 0.7180 | 0.4601 | 0.7267 | 0.8610 |
| CPC2 | ○ | 0.6780 | 0.9320 | 0.3840 | 0.4401 | 0.5439 | 0.7535 |
| FEELnc | ● | 0.7820 | 0.8730 | 0.6600 | 0.5816 | 0.7517 | 0.8674 |
| lncRNAnet | ○ | 0.8130 | 0.7350 | 0.9780 | 0.6631 | 0.8390 | 0.8690 |
| mRNN | ○ | 0.8720 | 0.9366 | 0.7980 | 0.7520 | 0.8617 | 0.9309 |

**Table S3.** Number of transcripts belonging to each class in the datasets of five different species.

| | Coding | Non-coding |
|---|---|---|
| *M. musculus* | 10,638 | 12,251 |
| *D. rerio* | 2,344 | 1,528 |
| *D. melanogaster* | 3,680 | 3,556 |
| *C. elegans* | 3,551 | 9,470 |
| *A. thaliana* | 13,986 | 3,853 |

**Table S4.** Comparison of classification performance in datasets of five different species. MCC: Matthews correlation coefficient. PRC: area under the precision-recall curve

| | Accuracy | Precision | Recall | MCC | F1-score | PRC |
|---|---|---|---|---|---|---|
| *M. musculus* | | | | | | |
| RNAsamba | 0.9680 | 0.9498 | 0.9831 | 0.9363 | 0.9661 | 0.9946 |
| CPAT | 0.9662 | 0.9829 | 0.9437 | 0.9325 | 0.9629 | 0.9928 |
| CPC2 | 0.9572 | 0.9675 | 0.9395 | 0.9141 | 0.9533 | 0.9895 |
| FEELnc | 0.8871 | 0.8213 | 0.9710 | 0.7872 | 0.8899 | 0.9855 |
| lncRNAnet | 0.9543 | 0.9566 | 0.9446 | 0.9082 | 0.9506 | 0.9868 |
| mRNN | 0.9485 | 0.9332 | 0.9578 | 0.8970 | 0.9454 | 0.9659 |
| *D. rerio* | | | | | | |
| RNAsamba | 0.9032 | 0.8534 | 0.9936 | 0.8153 | 0.9182 | 0.9657 |
| CPAT | 0.9315 | 0.9328 | 0.9553 | 0.8564 | 0.9439 | 0.9623 |
| CPC2 | 0.9367 | 0.9408 | 0.9556 | 0.8672 | 0.9481 | 0.9724 |
| FEELnc | 0.8692 | 0.8321 | 0.9829 | 0.7336 | 0.9012 | 0.9624 |
| lncRNAnet | 0.8970 | 0.8812 | 0.9590 | 0.7843 | 0.9185 | 0.9401 |
| mRNN | 0.8846 | 0.8644 | 0.9599 | 0.7590 | 0.9096 | 0.8607 |
| *D. melanogaster* | | | | | | |
| RNAsamba | 0.8320 | 0.7549 | 0.9916 | 0.6989 | 0.8572 | 0.9858 |
| CPAT | 0.9667 | 0.9661 | 0.9685 | 0.9334 | 0.9673 | 0.9919 |
| CPC2 | 0.9415 | 0.9379 | 0.9478 | 0.8831 | 0.9428 | 0.9837 |
| FEELnc | 0.8725 | 0.8098 | 0.9848 | 0.7624 | 0.8888 | 0.9903 |
| lncRNAnet | 0.8960 | 0.8425 | 0.9785 | 0.8024 | 0.9054 | 0.9857 |
| mRNN | 0.8495 | 0.7773 | 0.9867 | 0.7255 | 0.8696 | 0.9346 |
| *C. elegans* | | | | | | |
| RNAsamba | 0.9710 | 0.9075 | 0.9949 | 0.9309 | 0.9492 | 0.9983 |
| CPAT | 0.9815 | 0.9934 | 0.9383 | 0.9532 | 0.9651 | 0.9957 |
| CPC2 | 0.9907 | 0.9994 | 0.9665 | 0.9766 | 0.9827 | 0.9958 |
| FEELnc | 0.7967 | 0.5956 | 0.9992 | 0.6497 | 0.7463 | 0.9890 |
| lncRNAnet | 0.9478 | 0.8438 | 0.9918 | 0.8808 | 0.9118 | 0.9955 |
| mRNN | 0.9323 | 0.8048 | 0.9927 | 0.8506 | 0.8889 | 0.9937 |
| *A. thaliana* | | | | | | |
| RNAsamba | 0.9793 | 0.9798 | 0.9941 | 0.9383 | 0.9869 | 0.9995 |
| CPAT | 0.9409 | 0.9993 | 0.9252 | 0.8514 | 0.9608 | 0.9988 |
| CPC2 | 0.9635 | 0.9993 | 0.9541 | 0.9027 | 0.9762 | 0.9992 |
| FEELnc | 0.9466 | 0.9527 | 0.9812 | 0.8332 | 0.9667 | 0.9973 |
| lncRNAnet | 0.9663 | 0.9819 | 0.9750 | 0.9014 | 0.9784 | 0.9982 |
| mRNN | 0.9581 | 0.9764 | 0.9700 | 0.8774 | 0.9732 | 0.9951 |

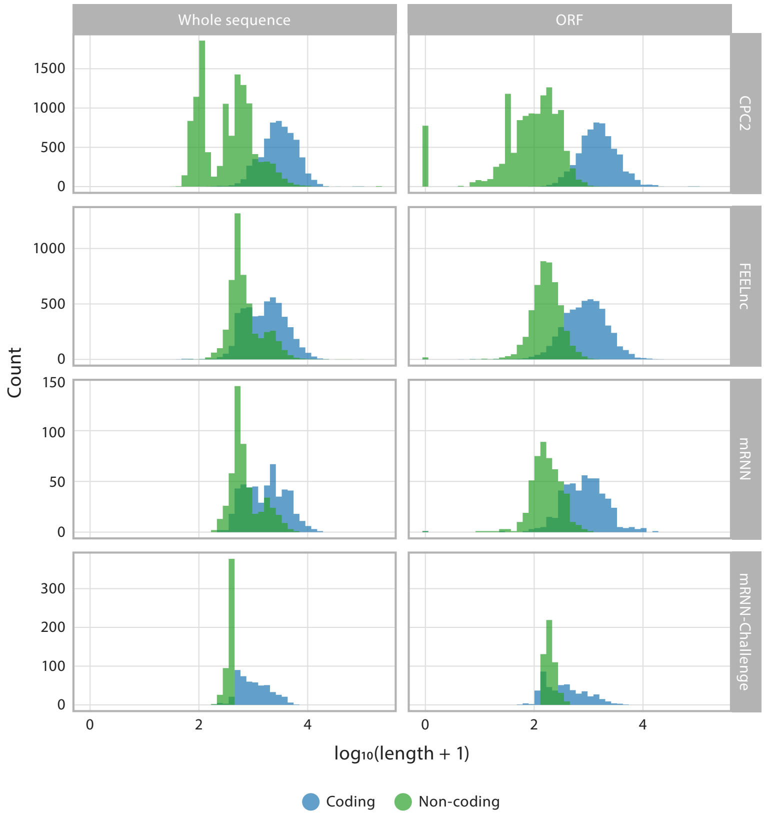


**Figure S1.** Length distribution of coding and non-coding transcripts and their extracted ORFs in each of the human test sets.

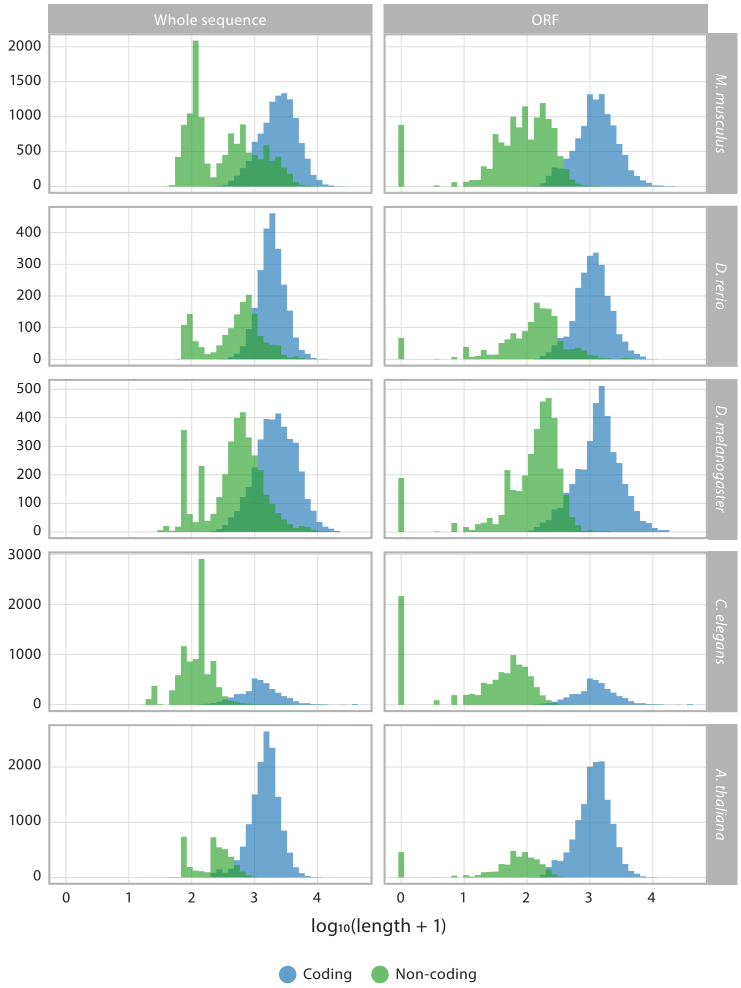


**Figure S2.** Length distribution of coding and non-coding transcripts and their extracted ORFs in each of five species datasets.

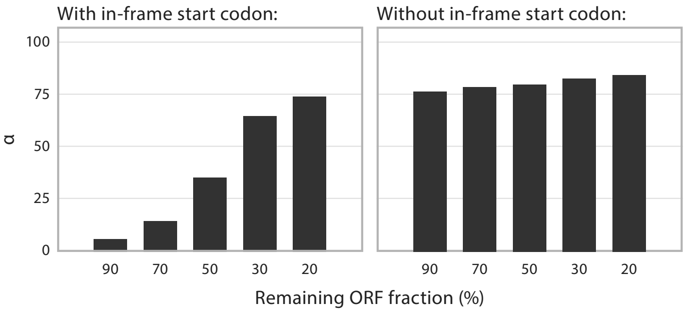


**Figure S3.** The median value of the α parameter increases as ORF-derived information diminishes.

**A.**

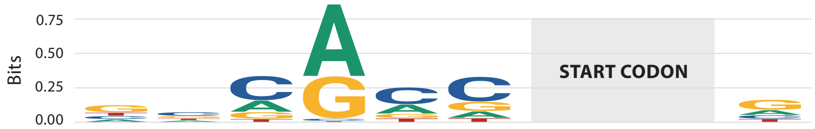


| | | | | | | | | | | |
|---|---|---|---|---|---|---|---|---|---|---|
| A | 0.17 | 0.18 | 0.25 | 0.61 | 0.27 | 0.15 | 1.00 | 0.00 | 0.00 | 0.23 |
| C | 0.19 | 0.39 | 0.53 | 0.02 | 0.49 | 0.55 | 0.00 | 0.00 | 0.00 | 0.16 |
| G | 0.44 | 0.23 | 0.15 | 0.36 | 0.13 | 0.21 | 0.00 | 0.00 | 1.00 | 0.46 |
| T | 0.20 | 0.20 | 0.07 | 0.01 | 0.11 | 0.09 | 0.00 | 1.00 | 0.00 | 0.15 |

**B.**

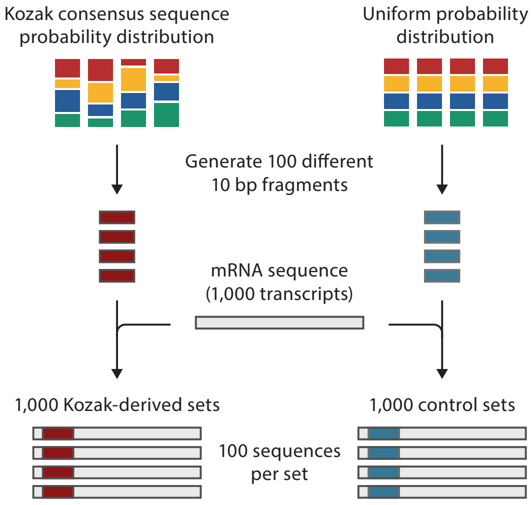


**Figure S4.** (A) Sequence logo and probability distribution of the Kozak consensus sequence. (B) Kozak sequence consensus probability distribution and a uniform probability distribution were used to generate two sets of sequences, each containing 100 fragments. Then, for each of 1,000 mouse mRNAs, the region spawning the positions -6 to +1 was exchanged by the fragments generated in the previous step, producing new transcripts containing both biological and computer-generated sequences.

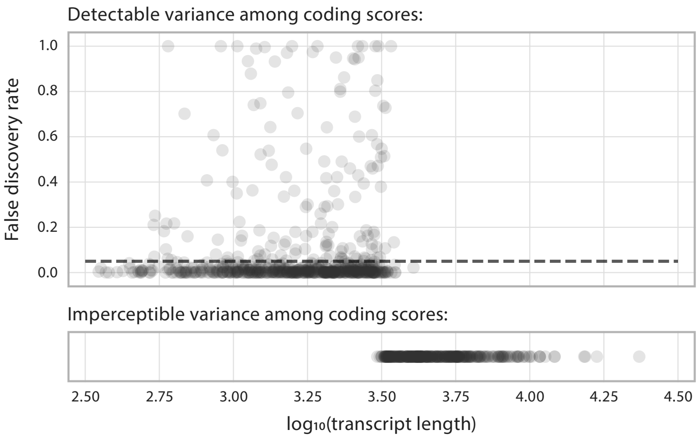


**Figure S5.** In most of the tested transcripts, the coding score of the sequences containing Kozak-derived fragments was significantly higher then the ones of sequences from the control set (FDR-adjusted $p\text{-}value \leq 0.05$, represented by the dashed line). No variation whatsoever could be detected among the coding scores of sequences longer than approximately 3,160 bp.

### Datasets

#### 1. CPC2

***Train:***

- http://cpc2.cbi.pku.edu.cn/download_file.php?filename=CPC2_coding_training_dataset.fa
- http://cpc2.cbi.pku.edu.cn/download_file.php?filename=CPC2_noncoding_training_dataset.fa

***Test (Human):***

- http://cpc2.cbi.pku.edu.cn/download_file.php?filename=mRNA_human.faa
- http://cpc2.cbi.pku.edu.cn/download_file.php?filename=lncRNA_human.faa
- http://cpc2.cbi.pku.edu.cn/download_file.php?filename=sncRNA_human.faa

***Test (M. musculus):***

- http://cpc2.cbi.pku.edu.cn/download_file.php?filename=mRNA_mouse.faa
- http://cpc2.cbi.pku.edu.cn/download_file.php?filename=lncRNA_mouse.faa
- http://cpc2.cbi.pku.edu.cn/download_file.php?filename=sncRNA_mouse.faa

***Test (D. rerio):***

- http://cpc2.cbi.pku.edu.cn/download_file.php?filename=mRNA_zebrafish.faa
- http://cpc2.cbi.pku.edu.cn/download_file.php?filename=lncRNA_zebrafish.faa
- http://cpc2.cbi.pku.edu.cn/download_file.php?filename=sncRNA_zebrafish.faa

***Test (D. melanogaster):***

- http://cpc2.cbi.pku.edu.cn/download_file.php?filename=mRNA_fruitfly.faa
- http://cpc2.cbi.pku.edu.cn/download_file.php?filename=lncRNA_fruitfly.faa
- http://cpc2.cbi.pku.edu.cn/download_file.php?filename=sncRNA_fruitfly.faa

***Test (C. elegans):***

- http://cpc2.cbi.pku.edu.cn/download_file.php?filename=mRNA_worm.faa
- http://cpc2.cbi.pku.edu.cn/download_file.php?filename=lncRNA_worm.faa
- http://cpc2.cbi.pku.edu.cn/download_file.php?filename=sncRNA_worm.faa

***Test (A. thaliana):***

- http://cpc2.cbi.pku.edu.cn/download_file.php?filename=mRNA_arabidopsis.faa
- http://cpc2.cbi.pku.edu.cn/download_file.php?filename=lncRNA_arabidopsis.faa
- http://cpc2.cbi.pku.edu.cn/download_file.php?filename=sncRNA_arabidopsis.faa

#### 2. FEELnc

***Train:***

- http://tools.genouest.org/data/tderrien/FEELnc_article_supplementary/human/Homo_sapiens.GRCh38.83_protein_coding_learning5k.fa
- http://tools.genouest.org/data/tderrien/FEELnc_article_supplementary/human/Homo_sapiens.GRCh38.83_antisense.lincRNA_learning5k.fa

***Test:***

- http://tools.genouest.org/data/tderrien/FEELnc_article_supplementary/human/Homo_sapiens.GRCh38.83_all_testing10k.fa
- http://tools.genouest.org/data/tderrien/FEELnc_article_supplementary/human/Homo_sapiens.GRCh38.83_all_testing10k.label

### 3. mRNN

***Train:***

- https://osf.io/pyqw9/download
- https://osf.io/afg3q/download

***Test:***

- https://osf.io/7utwj/download
- https://osf.io/v4xtb/download

***Test (Challenge set):***

- https://osf.io/njfb3/download
- https://osf.io/vfsu3/download
